# Identification and analysis of integrons and cassette arrays in bacterial genomes

**DOI:** 10.1101/030866

**Authors:** Jean Cury, Thomas Jové, Marie Touchon, Bertrand Néron, Eduardo PC Rocha

## Abstract

Integrons recombine gene arrays and favor the spread of antibiotic resistance. Their broader roles in bacterial adaptation remain mysterious, partly due to lack of computational tools. We made a program – IntegronFinder – to identify integrons with high accuracy and sensitivity. IntegronFinder is available as a standalone program and as a web application. It searches for *attC* sites using covariance models, for integron-integrases using HMM profiles, and for other features (promoters, *attI* site) using pattern matching. We searched for integrons, integron-integrases lacking *attC* sites, and clusters of *attC sites* lacking a neighboring integron-integrase in bacterial genomes. All these elements are especially frequent in genomes of intermediate size. They are missing in some key phyla, such as α-Proteobacteria, which might reflect selection against cell lineages that acquire integrons. The similarity between *attC* sites is proportional to the number of cassettes in the integron, and is particularly low in clusters of *attC* sites lacking integron-integrases. The latter are unexpectedly abundant in genomes lacking integron-integrases or their remains, and have a large novel pool of cassettes lacking homologs in the databases. They might represent an evolutionary step between the acquisition of genes within integrons and their stabilization in the new genome.

## Introduction

Integrons are gene-capturing platforms playing a major role in the spread of antibiotic resistance genes (reviewed in (1–4)). They have two main components (Figure 1). The first is made of the integron-integrase gene (*intI*)and its promoter (P_intl_), an integration site named *attI* (attachment site of the integron) and a constitutive promoter (Pc) for the gene cassettes integrated at the *attI* site (5). The second component is a cluster with up to 200 gene cassettes (6), most frequently transcribed in the opposite direction relative to the integron-integrase (7). Typical gene cassettes have an open reading frame (ORF) surrounded by *attC* recombination sites (attachment site of the cassette), but the presence of the ORF is not mandatory. Cassettes carrying their own promoters are expressed independently of Pc (8).

**Figure 1.**
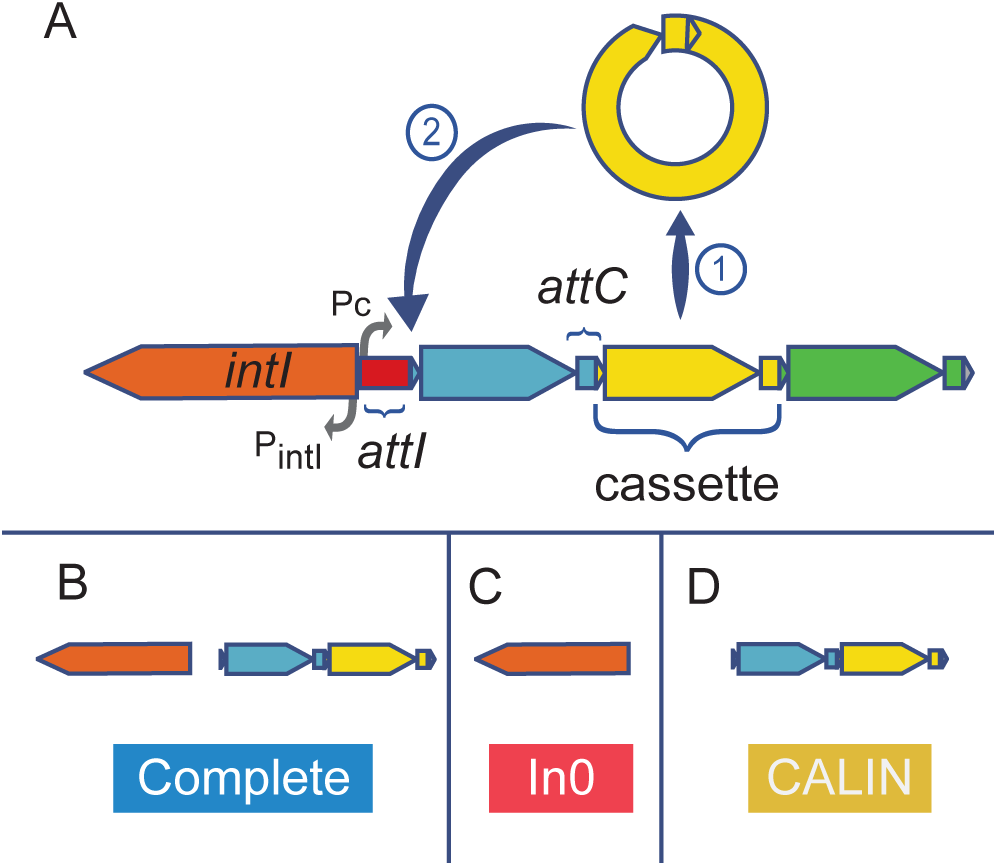
Schema of an integron and the three types of elements detected by IntegronFinder. (A) The integron is composed of a specific integron integrase gene (*intI*, orange), an *attI* recombination site (red), and an array of gene cassettes (blue, yellow and green). A cassette is typically composed of an ORF flanked by two *attC* recombination sites. The integron integrase has its own promoter (P_intI_). There is one constitutive promoter (Pc) for the cluster of cassettes. Cassettes rarely contain promoters. The integrase can excise a cassette (1) and/or integrate it at the *attI* site (2). (B) Complete integrons include an integrase and at least one *attC* site. (C) The In0 elements are composed of an integron integrase and no *attC* sites. (D) The <u>c</u>lusters of *<u>a</u>ttC* sites <u>l</u>acking <u>in</u>tegron-integrases (CALIN) are composed of at least two *attC* sites.

Integrase-mediated recombination between two adjacent *attC* sites leads to the excision of a circular DNA fragment composed of an ORF and an *attC* site. The recombination of the *attC* site of this circular DNA fragment with an *attI* site leads to integration of the fragment at the location of the latter (9,10). Integrons can use this mechanism to capture cassettes from other integrons or to rearrange the order of their cassettes (11). The mechanism responsible for the creation of new cassettes is unknown.

The most distinctive features of the integron are thus the integron-integrase gene (*intI)* and a cluster of *attC* sites (Figure 1). The integron-integrase is a site-specific tyrosine recombinase closely related to Xer proteins (12). Contrary to most other tyrosine recombinases, IntI recombines nucleotide sequences of low similarity (13,14), by recognizing specific structural features of the *attC* site (15,16) (Figure 2A). This is partly caused by the presence of a ~35 residues domain near the patch III region of the integron-integrase that is lacking in the other tyrosine recombinases (17). The integration of the *attC* site at *attI* produces chimeric *attI/attC* sites on one side and chimeric *attC/attC* sites on the other side of the cassette. This results in a cluster of chimerical *attC* sites with similar palindromic structures.

**Figure 2.**
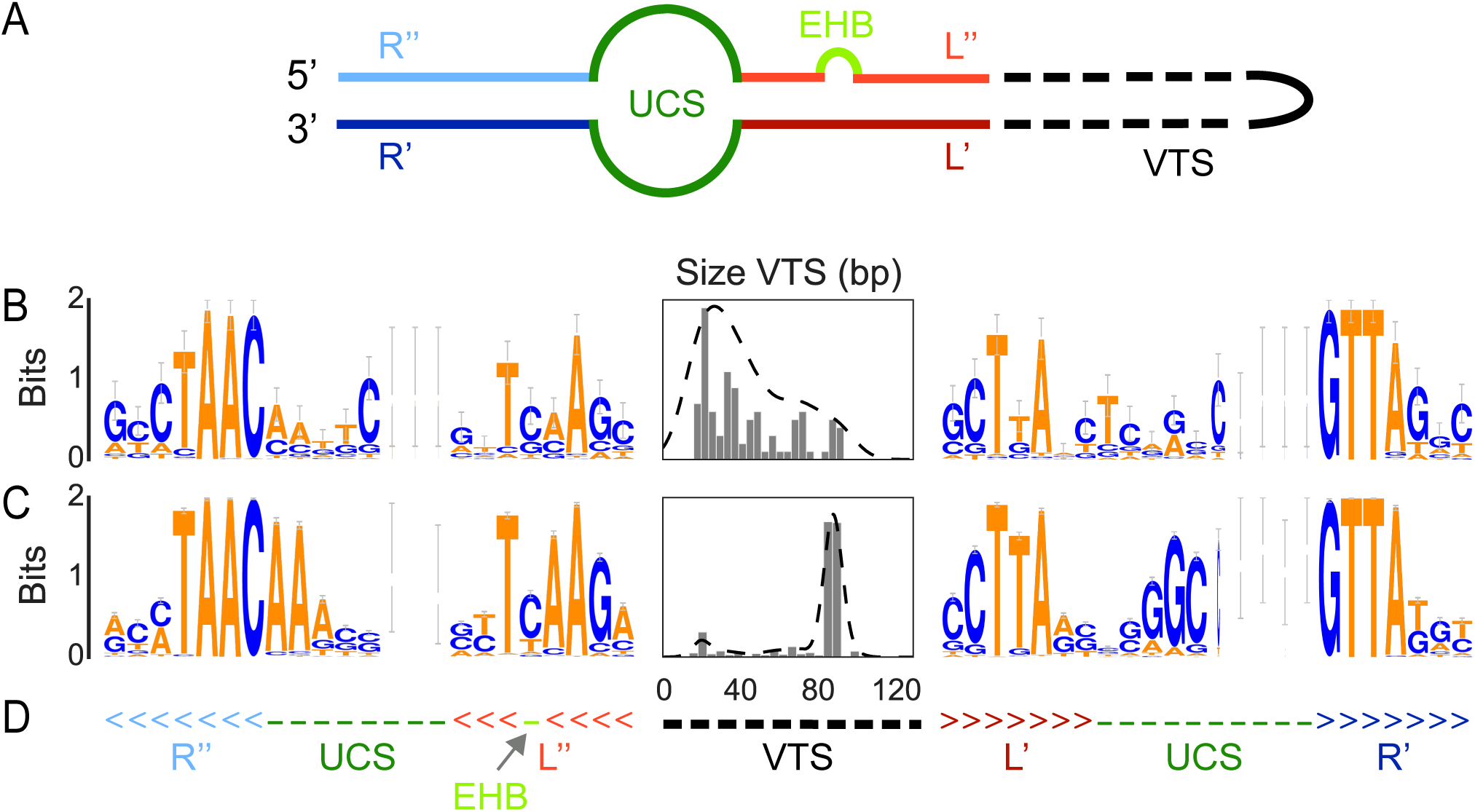
Characteristics of the *attC* sites. (A) Scheme of the secondary structure of a folded *attC* site. EHB stands for Extra Helical Bases. (B) Analysis of the *attC* sites used to build the model, including the WebLogo (70) of the R and L box and unpaired central spacers (UCS) and the histogram (and kernel density estimation) of the size of the variable terminal structure (VTS). The Weblogo represents the information contained in a column of a multiple sequence alignment (using the log2 transformation). The taller the letter is, the more conserved is the character at that position. The width of each column of symbols takes into account the presence of gaps. Thin columns are mostly composed of gaps. (C) Same as (B) but with the set of *attC* sites identified in complete integrons found in complete bacterial genomes. (D) Secondary structure used in the model in WUSS format, colors match those of (A).

Previous literature focused on integrons carrying antibiotic resistance genes. These integrons are often mobile, due to their association with transposons, and carry few cassettes (18,19,20). Most of the so-called mobile integrons can be classed in five classes, numbered 1 to 5. The IntI within each class show little genetic diversity, indicating their recent emergence from a much larger and diverse pool of integrons (7,21–23), possibly chromosomally encoded (24). For example, prototypical class 1 integrons were found on many chromosomes of non-pathogenic soil and freshwater β-Proteobacteria (25). By contrast, so-called chromosomal integrons are found in most strains of a species and carry cassettes encoding a wide range of functions. For example, the *Vibrio* spp. chromosomal integrons (initially called super-integrons due to their large number of cassettes (19)) encode virulence factors, secreted proteins, and toxin-antitoxin modules (21). The high similarity of *attC* sites within chromosomal, but not mobile, integrons of *Vibrio spp* has prompted the hypothesis that cassettes are created by chromosomal and spread by mobile integrons (26). The dichotomy between chromosomal and mobile integrons has been criticized (7) because integrons encoded in the chromosome may be in mobile elements (26) and/or have small arrays of cassettes (27).

The analysis of metagenomics data has unraveled a vast pool of novel cassettes in microbial communities (28). Although antibiotic-resistance integrons were found to be abundant in human-associated environments such as sewage (29–31), most cassettes in environmental datasets encode different functions (or genes of unknown function) (29–31). The study of these functions, and of the adaptive impact of integrons, has been hindered by the difficulty in identifying integron cassettes. The bottleneck in these analyses is the recognition of *attC* sites, for which few tools were made available. The program XXR identifies *attC* sites in the large *Vibrio* integrons using pattern-matching techniques (21). The programs ACID (6) (no longer available) and ATTACCA (32) (now a part of RAC, available under private login) were designed to search for class 1 to class 3 mobile integrons. Such classical motif detection tools based on sequence conservation identify *attC* sites only within restricted classes of integrons. They are inadequate to identify or align distantly related *attC* sites because their sequences are too dissimilar. Yet, these sites have conserved structural constraints that can be used to identify highly divergent sequences.

We built a program named IntegronFinder (Figure 3, https://github.com/gem-pasteur/Integron_Finder) to detect integrons and their most distinctive components: the integron-integrase with the use of HMM profiles and the *attC* sites with the use of a covariance model (Figure 2). Covariance models use stochastic context-free grammars to model the constraints imposed by sequence pairing to form secondary structures. Such models have been previously used to detect structured motifs, such as tRNAs (33). They provide a good balance between sensitivity, the ability to identify true elements even if very diverse in sequence, and specificity, the ability to exclude false elements (34). They are ideally suited to model elements with high conservation of structure and poor conservation of sequence, such as *attC* sites. IntegronFinder also annotates known *attI* sites, P_intI_ and P_C_, and any pre-defined type of protein coding genes in the cassettes (*e.g.*, antibiotic resistance genes). IntegronFinder was built to accurately identify integron-integrases and *attC* sites of any generic integron. Importantly, we provide the program on a webserver that is free, requires no login, and has a long track record of stability (35) (http://mobyle.pasteur.fr/cgi-bin/portal.py#forms::integron_finder). We also provide a standalone application for large-scale genomics and metagenomics projects. We used IntegronFinder to identify integrons in bacterial genomes and to characterize their distribution and diversity.

**Figure 3.**
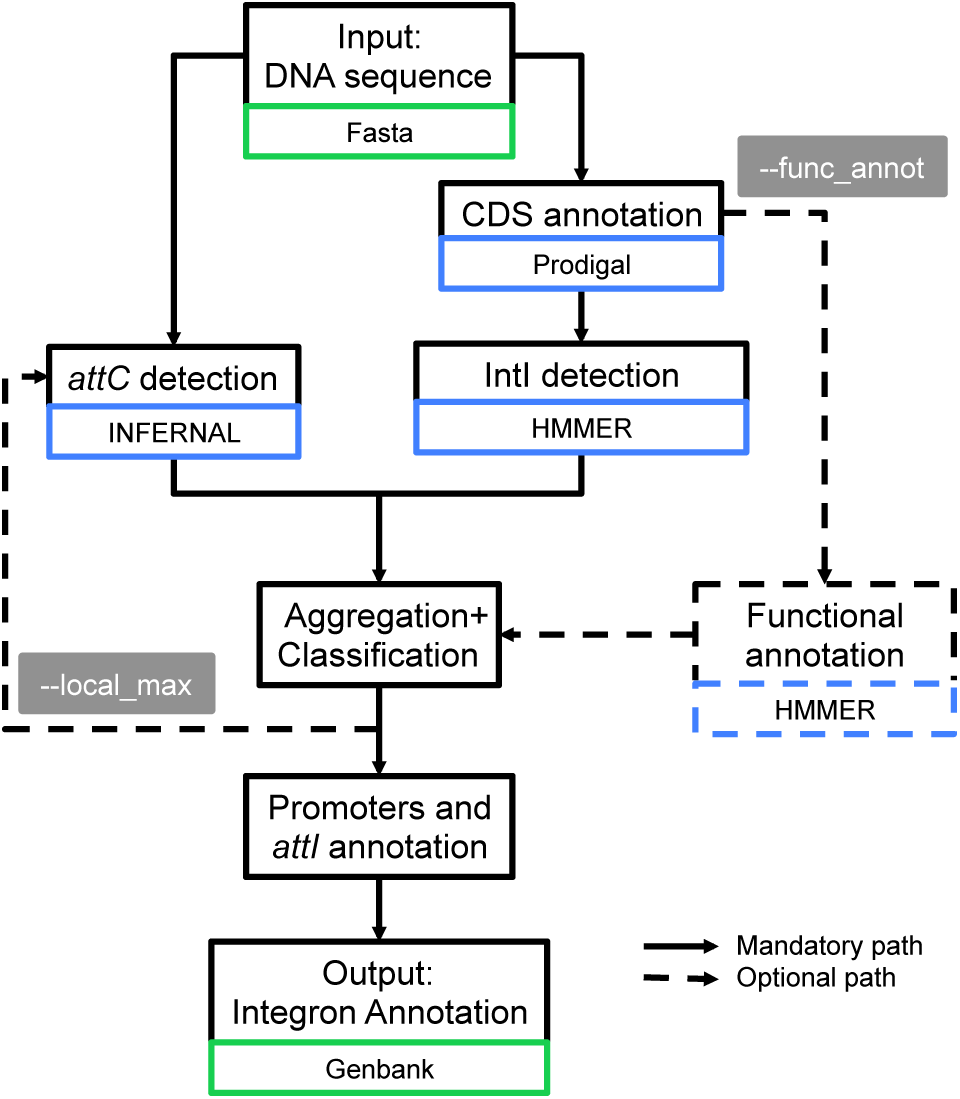
Diagram describing the different steps used by IntegronFinder to identify and annotate integrons. Solid lines represent the default mode, dotted lines optional modes. Blue boxes indicate the main dependency used for a given step. Green boxes indicate the format of the file needed for a given step.

## Material and Methods

### Data

The sequences and annotations of complete genomes were downloaded from NCBI RefSeq (last accessed in November 2013, ftp://ftp.ncbi.nih.gov/genomes/Bacteria/). Our analysis included 2484 bacterial genomes (see Table S1). We used the classification of replicons in plasmids and chromosomes as provided in the GenBank files. Our dataset included 2626 replicons labeled as chromosomes and 2006 as plasmids. The *attC* sites used to build the covariance model and the accession numbers of the replicons manually curated for the presence or absence of *attC* sites were retrieved from INTEGRALL, the reference database of integron sequences (http://integrall.bio.ua.pt/) (36). We used a set of 291 *attC* sites (Supplementary File 1) to build and test the model, and a set of 346 sequences with expert annotation of 596 *attC* sites to analyze the quality of the program predictions (Table S2a and S2b).

### Protein profiles

We built a protein profile for the region specific to the integron tyrosine recombinase. For this, we retrieved the 402 IntI homologues from the Supplementary file 11 of Cambray et al (37). These proteins were clustered using uclust 3.0.617 (38) with a threshold of 90% identity to remove very closely related proteins (the largest homologs were kept in each case). The resulting 79 proteins were used to make a multiple alignment using MAFFT (39) (‐‐globalpair ‐‐maxiterate 1000). The position of the specific region of the integron-integrase in *V. cholerae* was mapped on the multiple alignments using the coordinates of the specific region taken from (17). We recovered this section of the multiple alignment to produce a protein profile with hmmbuild from the HMMer suite version 3.1b1 (40). This profile was named intI_Cterm (Supplementary File 2).

We used 119 protein profiles of the Resfams database (core version, last accessed on January 20, 2015 v1.1), to search for genes conferring resistance to antibiotics (http://www.dantaslab.org/resfams (41)). We retrieved from PFAM the generic protein profile for the tyrosine recombinases (PF00589, phage_integrase, http://pfam.xfam.org/(42)). All the protein profiles were searched using hmmsearch from the HMMer suite version 3.1b1. Hits were regarded as significant when their e-value was smaller than 0.001 and their alignment covered at least 50% of the profile.

### Construction and analysis of *attC* models

We built a covariance model for the *attC* sites (Supplementary File 3). These models score a combination of sequence and secondary structure consensus (33) (with the limitation that these are DNA not RNA structures). To produce the *attC* models, 96 *attC* sites (33%) were chosen randomly from 291 known *attC* (see Data). The alignments were manually curated to keep the known conserved regions of the R and L boxes aligned in blocks. The unpaired central spacers (UCS) and the variable terminal structure (VTS) were not aligned because they were poorly conserved in sequence and length. Gaps were inserted in the middle of the VTS sequence as needed to keep the blocks of R and L boxes aligned. The consensus secondary structure was written in WUSS format beneath the aligned sequences (Supplementary File 4). The model was then built with INFERNAL 1.1 (34) using *cmbuild* with the option "‐‐hand". This option allows the user to set the columns of the alignment that are actual matches (consensus). This is crucial for the quality of the model, because most of the columns in the R and L boxes would otherwise be automatically assigned as inserts due to the lack of sequence conservation. The R-UCS-L sections of the alignment were chosen as the consensus region, and the VTS was designed as a gap region. We used *cmcalibrate* from INFERNAL 1.1 to fit the exponential tail of the covariance model e-values, with default options. The model was used to identify *attC* sites using INFERNAL with two alternative modes. The default mode uses heuristics to reduce the sequence space of the search. The Inside algorithm is more accurate, but computationally much more expensive (typically 10^4^ times slower) (34). By default, the *attC* sites were kept for further analysis when their e-value were below 1. The user can set this value (option ‐‐evalue_attc).

### Identification of promoters and *attI* sites

The sequences of the Pc promoters, of the P_intI_ promoters, and of the *attI* site were retrieved from INTEGRALL for the integrons of class 1, 2 and 3 when available (see Table S3). We searched for exact matches of these sequences (accepting no indel nor mismatch) using pattern matching as implemented in the search function of the Bio.motifs module of Biopython v1.65 (43).

### Overview of IntegronFinder:a program for the identification of *attC* sites, *intl* genes, integrons, and CALIN elements

The input of IntegronFinder is a sequence of DNA in FASTA format. The sequence is annotated with Prodigal v2.6.2 (44) using the default mode for replicons larger than 200kb and the metagenomic mode for smaller replicons ("-p meta” in Prodigal) (Figure 3). In the present work, we omitted the annotation part and used the NCBI RefSeq annotations because they are curated. The annotation step is particularly useful to study newly acquired sequences or poorly annotated ones.

The program searches for the two protein profiles of the integron-integrase using hmmsearch with default parameters from HMMER suite version 3.1b1 and for the *attC* sites with the default mode of cmsearch from INFERNAL 1.1 (Figure 3). Two *attC* sites are put in the same cluster if they are less than 4 kb apart on the same strand. The clusters are built by transitivity: an *attC* site less than 4 kb from any *attC* site of a cluster is integrated in that cluster. Clusters are merged when localized less than 4 kb apart. The threshold of 4kb was determined empirically as a compromise between sensitivity (large values decrease the probability of missing cassettes) and specificity (small values are less likely to put together two independent integrons). More precisely, the threshold is twice the size of the largest known cassettes (~2 kb (6)). This guarantees that even in the worst case (largest known cassettes) two *attC* sites will be clustered if an intervening site was not detected. Importantly, the user can set this threshold (“‐‐distance_thresh” in IntegronFinder).

The results of the searches for the elements of the integron are put together to class the loci in three categories (Figure 1 – B, C, D). (1) The elements with *intI* and at least one *attC* site were named complete integrons. The word complete is meant to characterize the presence of both elements; we cannot ascertain the functionality or expression of the integron. (2) The *In0* elements have *intI* but no recognizable *attC sites*. We do not strictly follow the original definition of In0, which also includes the presence of an *attI* (45), because this sequence is not known for most integrons (and thus cannot be searched for). (3) The <u>c</u>luster of *<u>a</u>ttC* site <u>l</u>acking <u>in</u>tegron-integrase (CALIN) have at least two *attC* sites and lack nearby *intI*.

To obtain a better compromise between accuracy and running time, IntegronFinder can re-runs INFERNAL to search for *attC* sites with more precision using the Inside algorithm ("‐‐max" option in INFERNAL), but only around previously identified elements ("‐‐local_max" option in IntegronFinder). More precisely, if a locus contains an integron-integrase and *attC* sites (complete integron), the search is constrained to the strand encoding *attC* sites between the end of the integron-integrase and 4kb after its most distant *attC*. If other *attC* sites are found after this one, the search is extended by 4 kb in that direction until no more new sites are found. If the element contains only *attC* sites (CALIN), the search is performed on the same strand on both directions. If the integron is In0, the search for *attC* sites is done on both strands in the 4kb flanking the integron-integrase on each side. The program then searches for promoters and *attI* sites near the integron-integrase. Finally, it can annotate the integron genes’ cassettes (defined in the program as the CDS found between *intI* and 200 bp after the last *attC* site, or 200 bp before the first and 200 bp after the last *attC* site if there is no integron-integrase) using a database of protein profiles (option "‐‐func_annot"). For example, in the present study we used the ResFams database to search for antibiotic resistance genes. One can use any hmmer-compatible profile databases with the program.

The program outputs tabular and GenBank files listing all the identified genetic elements associated with an integron. The program also produces a figure in pdf format representing each complete integron. For an interactive view of all the hits, one can use the GenBank file as input in specific programs such as Geneious (46).

The user can change the profiles of the integrases and the covariance model of the *attC* site. Thus, if novel models of *attC* sites were to be built in the future, *e.g.*, for novel types of *attC* sites, they could easily be plugged in IntegronFinder.

### Phylogenetic analyses

We have made two phylogenetic analyses. One analysis encompasses the set of all tyrosine recombinases and the other focuses on IntI. The phylogenetic tree of tyrosine recombinases (Figure S1) was built using 204 proteins, including: 21 integrases adjacent to *attC* sites and matching the PF00589 profile but lacking the intI_Cterm domain, seven proteins identified by both profiles and representative of the diversity of IntI, and 176 known tyrosine recombinases from phages and from the literature (12). We aligned the protein sequences with Muscle v3.8.31 with default options (47). We curated the alignment with BMGE using default options (48). The tree was then built with IQ-TREE multicore version 1.2.3 with the model LG+I+G4. This model was the one minimizing the Bayesian Information Criterion (BIC) among all models available ("‐‐m TEST" option in IQ-TREE). We made 10000 ultra fast bootstraps to evaluate node support (Figure S1, Tree S1).

The phylogenetic analysis of IntI was done using the sequences from complete integrons or In0 elements (*i.e.*, integrases identified by both HMM profiles) (Figure S2). We added to this dataset some of the known integron-integrases of class 1, 2, 3, 4 and 5 retrieved from INTEGRALL. Given the previous phylogenetic analysis we used known XerC and XerD proteins to root the tree. Alignment and phylogenetic reconstruction were done using the same procedure; except that we built ten trees independently, and picked the one with best log-likelihood for the analysis (as recommended by the IQ-TREE authors (49)). The robustness of the branches was assessed using 1000 bootstraps (Figure S2, Tree S2, Table S4).

### Pan-genomes

Pan-genomes are the full complement of genes in the species. They were built by clustering homologous proteins into families for each of the species (as previously described in (50)). Briefly, we determined the lists of putative homologs between pairs of genomes with BLASTP (default parameters) and used the e-values (<10^−4^) to cluster them using SILIX (51). SILIX parameters were set such that a protein was homologous to another in a given family if the aligned part had at least 80% identity and if it included more than 80% of the smallest protein. We built pan-genomes for the 12 species having at least 4 complete genomes available in Genbank RefSeq and encoding at least one IntI. The genomes of these species carried 40% of the complete integrons in our dataset. We did not build a pan-genome for *Xanthomonas oryzae* because it contained too many rearrangements and repeated elements (52).

For a given species we computed the pattern of presence and absence of each integron-integrase protein family and the frequency of the integron-integrase within the species.

### Integron classification

We used two criteria to class integrons: frequency within the species’ genomes (Figure S3) and number of cassettes (Figure S4). Integrons were classed as sedentary chromosomal integrons (as named by (4)) when their frequency in the pan-genome was 100%, or when they contained more than 19 *attC* sites. They were classed as mobile integrons when missing in more than 40% of the species’ genomes, when present on a plasmid, or when the integron-integrase was from classes 1 to 5. The remaining integrons were classed as “other”.

### Pseudo-genes detection

We translated the six reading frames of the region containing the CALIN elements (10kb on each side) to detect *intI* pseudo-genes. We then ran hmmsearch with default options from HMMER suite v3.1b1 to search for hits matching the profile intI_Cterm and the profile PF00589 among the translated reading frames. We recovered the hits with e-values lower than 10^−3^ and alignments covering more than 50% of the profiles.

### IS detection

We identified insertion sequences (IS) by searching for sequence similarity between the genes present 4kb around or within each genetic element and a database of IS from ISFinder (53). Details can be found in (54).

### Detection of cassettes in INTEGRALL

We searched for sequence similarity between all the CDS of CALIN elements and the INTEGRALL database using BLASTN from BLAST 2.2.30+. Cassettes were considered homologous to those of INTEGRALL when the BLASTN alignment showed more than 40% identity.

## Results

### Models for *attC* sites

We selected a manually curated set of 291 *attC* sites representative of the diversity of sequences available in INTEGRALL (see Methods). We randomly sampled a third of them to build a covariance model of the *attC* site and set aside the others for subsequent validation. The characteristics of these sequences were studied in detail (Figure 2A), notably concerning the R and L boxes, the UCS, and the EHB (15). The positions of the so-called Conserved Triplet (AAC and the complementary GTT) were more conserved than the others (Figure 2B and 2D). The length and sequence of the VTS were highly variable, between 20 and 100 nucleotides long, as previously observed (1).

We used the covariance model to search for *attC* sites on 2484 complete bacterial genomes. The genomic *attC* sites showed stronger consensus sequences and more homogeneous VTS lengths than those used to build the model (Figure 2C). The analyses of sensitivity in the next paragraph show that our model missed very few sites. Hence, the differences between the initial and the genomic *attC* sites might be due to our explicit option of using diverse sequences to build the model (to maximize diversity). They may also reflect differences between mobile integrons (very abundant in INTEGRALL) and integrons in sequenced bacterial genomes (where a sizeable fraction of cassettes were identified in *Vibrio* spp.).

We tested the ability of the covariance model to identify known sequences within pseudo-genomes built by randomizing dinucleotides from genomes with varying G+C content (Table S5). We integrated in each pseudo-genome five *attC* sites (among the 195 of the validation set) at 2 kb intervals. We searched for *attC* sites in these genomes and found very few false positives in both run modes (~0.03 FP/Mb, Figure 4, see Methods for details). The proportion of true *attC* sites actually identified (sensitivity), was 61 % for the default mode and 88% for the most accurate mode (with option "‐‐local_max"). We identified at least two of the *attC* sites in 99% of the clusters (with the most accurate mode). Hence, clusters could be identified even when some *attC* sites were missed. The sensitivity of the model showed very little dependency on genome G+C composition in all cases (Figure 4).

**Figure 4.**
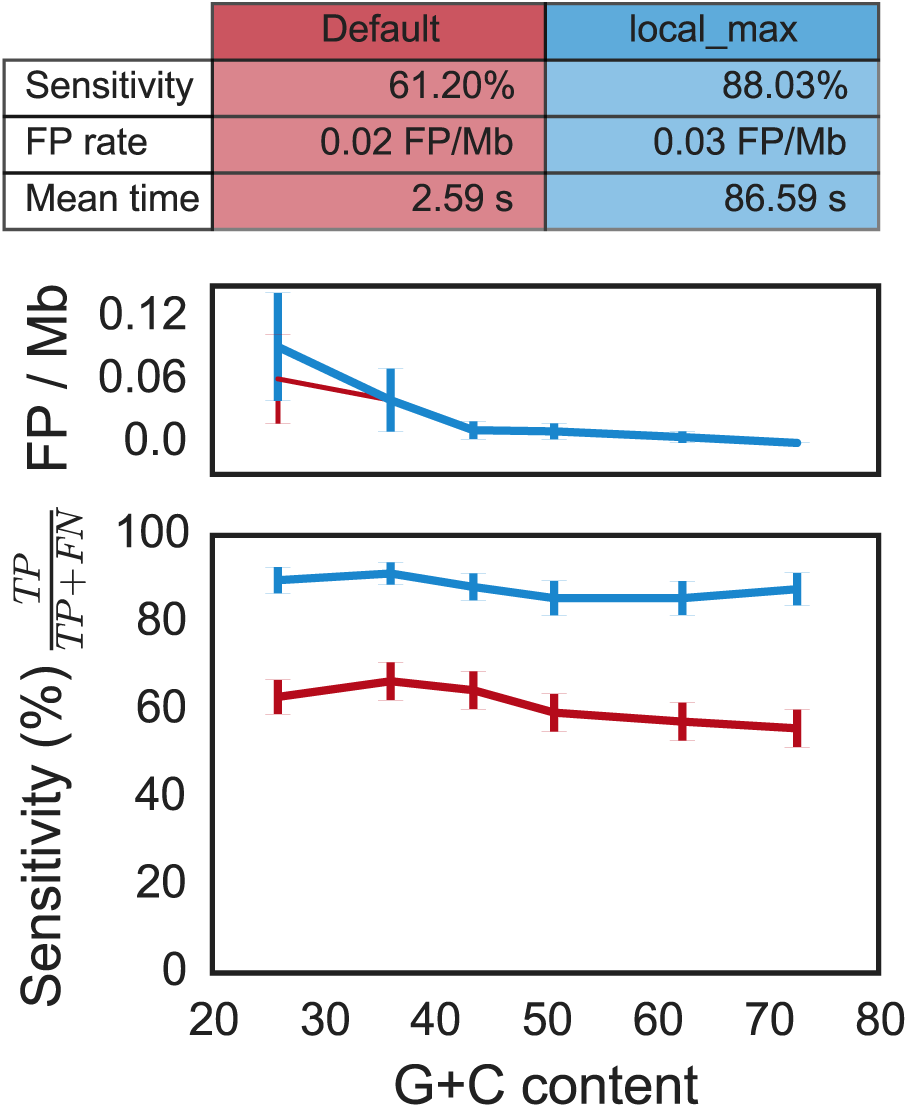
Quality assessment of the *attC* sites covariance model on pseudo-genomes with varying G+C content and depending on the run mode (default and "‐‐local_max"). (Top) Table resuming the results. The mean time is the average running time per pseudo-genome on a Mac Pro, 2 × 2.4 GHz 6–Core Intel Xeon, 16 Gb RAM, with options ‐‐cpu 20 and ‐‐no-proteins. (Middle) Rate of false positives per megabase (Mb) as function of the G+C content. (Bottom) Sensitivity (or true positive rate) as function of the G+C content. The red line depicts results obtained with the default parameters, and the blue line represents results obtained with the accurate parameters ("‐‐local_max" option). Vertical lines represent standard error of the mean. There is no correlation with G+C content (all spearman ρ ϵ [-0.12; −0.04] and all p-values > 0.06)

We then searched for *attC* sites in sequences annotated for the presence of integrons in INTEGRALL (Table S2a). The search was performed in 346 sequences containing 596 known *attC* sites. We found 570 *attC* sites with the most accurate mode (96% of sensitivity, Table S2b). We missed 26 of the known *attC* sites, among which 15 were on the integron edges, and were probably missed because of the absence of R’ box on the 3’ side. All the 57 sequences annotated as In0 in INTEGRALL also lacked *attC* sites in our analysis. We found 247 *attC* sites missing in the annotations of INTEGRALL. If *attC* annotations in INTEGRALL were perfect and all these sites were false, the rate of false positives of our analysis would be 0.72 per Mb. However, about 90% of these non-annotated *attC sites* were found in clusters of two *attC* sites or more, which suggests that they are real *attC* sites. If all isolated *attC* sites were false positives (and the only ones), then the false positive rate would be 0.07 FP/Mb, *i.e.*, less than one false *attC* site per genome.

These analyses showed a rate of false positives between 0.03 FP/Mb and 0.72 FP/Mb. The probability of having a cluster of two or more false *attC* sites by chance (within 4 kb) given this density of false positives is between 4.10^−6^ and 7.10^−9^ depending on the false positive rate (assuming a Poisson process). Hence, the clusters of *attC* sites given by our model are extremely unlikely to be false positives.

### Identification of integron-integrases

We identified tyrosine recombinases using the protein profile PF00589 (from PFAM). To distinguish IntI from the other tyrosine recombinases, we built an additional protein profile corresponding to the IntI specific region near the patch III domain (17) (henceforth named intI_Cterm, see Methods). We found 215 proteins matching both profiles in the complete genomes of bacteria. Only six genes matched intI_Cterm but not PF00589. There were 18,808 occurrences of PF00589 not matching intI_Cterm, among which only 50 co-localized with an *attC* site. Among the latter, 29 were in genomes that encoded IntI elsewhere in the replicon (Figure S5). The remaining 21 integrases were scattered in the phylogenetic tree of tyrosine recombinases, and only four of them were placed in an intermediate position between IntI and Xer (Figure S1). These four sequences resembled typical phage integrases at the region of the patch III domain characteristic of IntI and they colocalized with very few *attC* sites (always less than three). This analysis strongly suggests that tyrosine recombinases lacking the intI_Cterm domain identified near *attC* sites are not Intl.

Most *intI* genes identified in bacterial genomes co-localized with *attC* sites (76%, Figure S5). It is difficult to assess if the remaining *intI* genes are true or false, since In0 elements have often been described in the literature (45,55). We were able to identify IntI in the integrons of class 1 to class 5, as well as in well-known chromosomal integrons (*e.g.*, in *Vibrio* super-integrons). We also identified all In0 elements in the INTEGRALL dataset mentioned above. Overall, these results show that IntI could be identified accurately using the intersection of both protein profiles.

We built a phylogenetic tree of the 215 IntI proteins identified in genomes (Figure S2). Together with the analysis of the broader phylogenetic tree of tyrosine recombinases (Figure S1), this extends and confirms previous analyses (1,7,23,56): 1) The XerC and XerD sequences are close outgroups. 2) The IntI are monophyletic. 3)Within IntI, there are early splits, first for a clade including class 5 integrons, and then for *Vibrio* super-integrons. On the other hand, a group of integrons displaying an integron-integrase in the same orientation as the *attC* sites (inverted integron-integrase group) was previously described as a monophyletic group (7), but in our analysis it was clearly paraphyletic (Figure S2, column F). Notably, in addition to the previously identified inverted integron-integrase group of certain *Treponema* spp., a class 1 integron present in the genome of *Acinetobacter baumannii* 1656-2 had an inverted integron-integrase.

### Integrons in bacterial genomes

We built a program – IntegronFinder – to identify integrons in DNA sequences. This program searches for *intI* genes, *attC* sites, clusters them in function of their colocalization, and then annotates cassettes and other accessory genetic elements (see Figure 3 and Methods). The use of this program led to the identification of 215 IntI and 4597 *attC* sites in complete bacterial genomes. The combination of this data resulted in a dataset of 164 complete integrons, 51 In0, and 279 CALIN elements (see Figure 1 for their description). The observed abundance of complete integrons is compatible with previous data (7). While most genomes encoded a single integron-integrase, we found 36 genomes encoding more than one, suggesting that multiple integrons are relatively frequent (20% of genomes encoding integrons). Interestingly, while the literature on antibiotic resistance often reports the presence of integrons in plasmids, we only found 24 integrons with integron-integrase (20 complete integrons, 4 In0) among the 2006 plasmids of complete genomes. All but one of these integrons were of class 1 (96%).

The taxonomic distribution of integrons was very heterogeneous (Figure 5 and S6). Some clades contained many elements. The foremost clade was the γ-Proteobacteria among which 20% of the genomes encoded at least one complete integron. This is almost four times as much as expected given the average frequency of these elements (~6%, χ^2^ test in a contingency table, P < 0.001). The β-Proteobacteria also encoded numerous integrons (~10% of the genomes). In contrast, all the genomes of Firmicutes, Tenericutes, and Actinobacteria lacked complete integrons. Furthermore, all 243 genomes of α-Proteobacteria, the sister-clade of γ and β-Proteobacteria, were devoid of complete integrons, In0, and CALIN elements. Interestingly, much more distantly related bacteria such as Spirochaetes, Chlorobi, Chloroflexi, Verrucomicrobia, and Cyanobacteria encoded integrons (Figure 5 and Figure S6). The complete lack of integrons in one large phylum of Proteobacteria is thus very intriguing.

**Figure 5.**
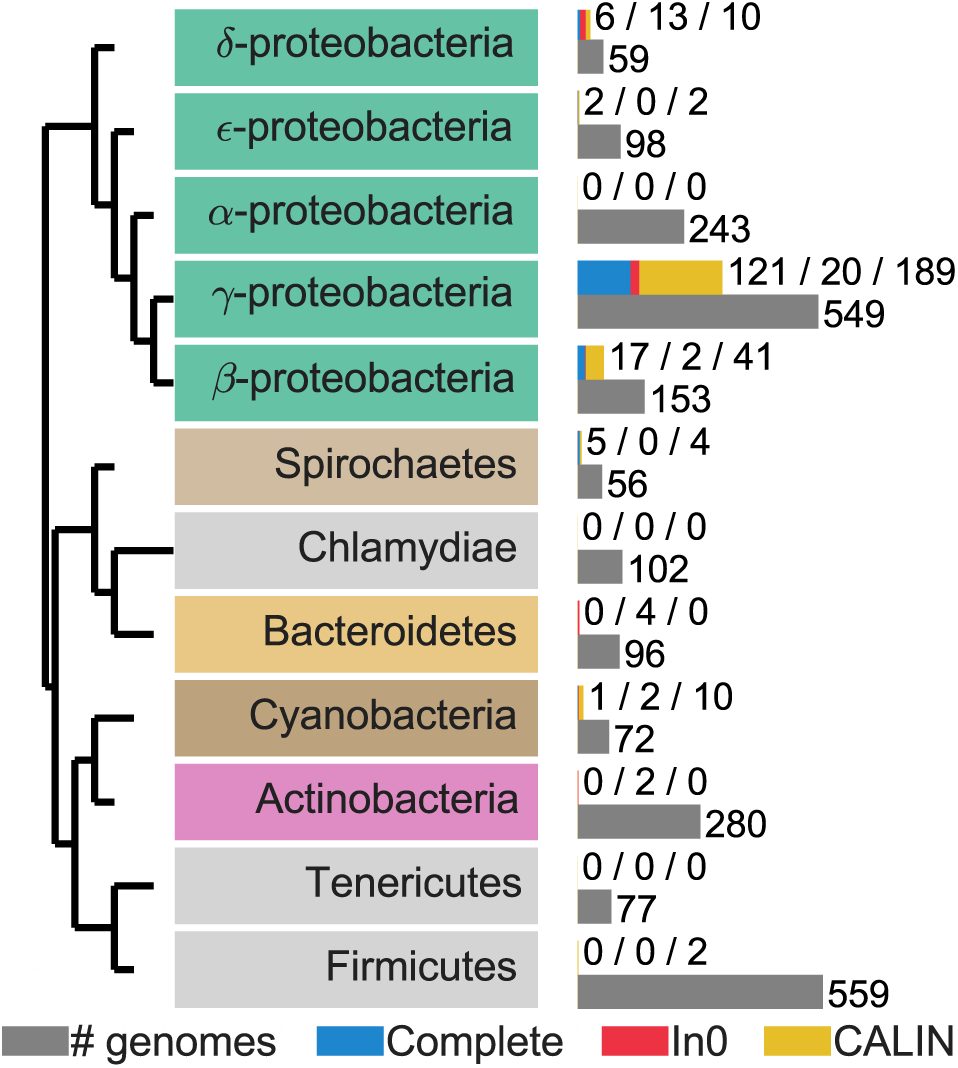
Taxonomic distribution of integrons in clades with more than 50 complete genomes sequenced. The grey bar represents the number of genomes sequenced for a given clade. The blue bar represents the number of complete integrons, the red bar the number of In0, and the yellow bar the number of CALIN. The colored text boxes refer to the colors in Figure S2.

We searched for genes encoding antibiotic resistance in integron cassettes (see Methods). We identified such genes in 105 cassettes, *i.e.*, in 3% of all cassettes from complete integrons (3116 cassettes). Most resistance cassettes were found in class 1 to 5 integrons (90% of them), even if the latter contained only 4.5% of all cassettes. This fits previous observations that integrons lacking antibiotic resistance determinants are very frequent in natural populations (25,28).

The association between genome size and the frequency of integrons has not been studied before. We binned the genomes in terms of their size and analyzed the frequency of complete integrons, In0, and CALIN. This showed a clearly nonmonotonic trend (Figure 6). This distribution was not homogeneous in the different size categories (χ^2^ test in a contingency table, p-values <1.10^−4^ for complete, CALIN and In0). The same result was observed in a complementary analysis using only integrons from Gamma-Proteobacteria (Figure S7). Very small genomes lack complete integrons, intermediate size genomes accumulate most of the integrons, and the largest genomes encode few. Importantly, the same trends were observed for In0 and CALIN. Hence, the frequency of integrons is maximal for genomes of intermediate size (4 to 6 Mb).

**Figure 6.**
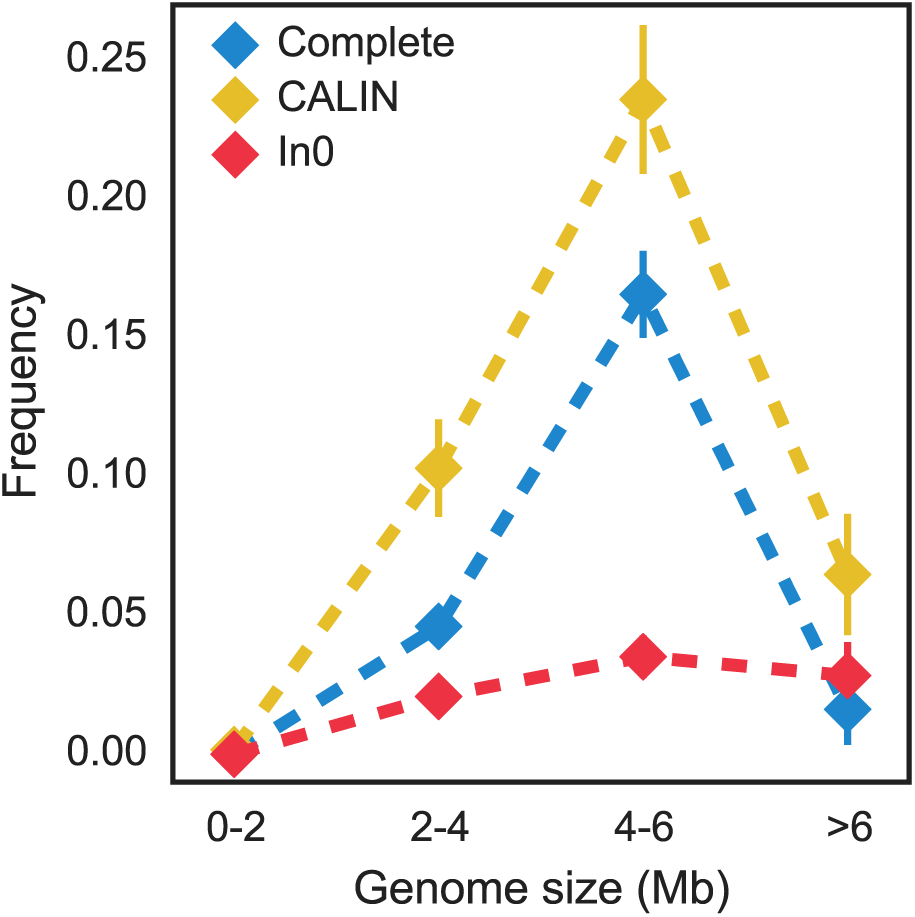
Frequency of integrons and related elements as a function of the genome size. Vertical bar represents standard error of the mean. The sample size in each bin is: 608 [0-2], 912 [2-4], 712 [4-6], and 247 [>6].

### Unexpected abundance of CALIN elements

The number of *attC* sites lacking nearby integron-integrases was unexpectedly high. We found 431 occurrences of isolated single *attC* sites among the 1879 *attC* sites lacking an integrase. If these sites were all false, and were the only false ones, then the observed rate of false positives can be estimated at 0.047 FP/Mb. This is within the range of the rates of false positives observed in the sensitivity analysis (between 0.02 FP/Mb and 0.72 FP/Mb). The probability that CALIN elements are false positives is exceedingly small for these rates of false positives. Therefore, we discarded single *attC* sites and kept the 279 clusters with two or more sites (CALIN) for the subsequent analyses. The CALIN resemble mobile integrons in terms of the number of cassettes: 83% had fewer than six *attC* sites and only 6.6% had more than 10 (Figure S8). Nevertheless, some few CALIN were very large, with up to 114 *attC* sites. Furthermore, their cassettes were remarkably different from those of mobile integrons: only 147 out of the 1933 cassettes were homologous to those reported in INTEGRALL and only 31 carried antibiotic resistance genes (to be compared with 70% among class 1 to class 5 integrons and with 0.4% among the other complete integrons). Hence, CALIN are relatively small on average (5 *attC* sites) but may contain several tens of *attC* sites, and have many previously unknown gene cassettes.

The CALIN elements might have arisen from the loss of the integrase in a previously complete integron. Therefore, we searched for pseudogenes matching the specific IntI_Cterm domain less than 10kb away from CALIN. We found such pseudo-genes near 15 out of 279 CALIN elements. It is worth noting that out of the 15 hits, 11 pseudo-genes were also matched by the PF00589 profile, which is consistent with the idea that they previously encoded IntI. Overall, our analysis showed that most CALIN (95%) are not close to recognizable *intI* pseudogenes.

We enquired on the possibility that some CALIN might actually be part of an integron and that we have missed this association because of the small 4 kb threshold used in the definition of the clusters. To test this hypothesis, we re-run IntegronFinder with a distance threshold of 10kb. This analysis found 252 of the previously identified 279 CALIN elements, the remaining 27 being merged with a integron or In0 element with the 10 kb threshold. This shows that increasing the distance threshold in the clustering procedure does not significantly change the observed abundance of CALIN.

Chromosomal rearrangements (integrations, translocations, or inversions) may split integrons and separate some cassettes from the neighborhood of the integron-integrase, thus producing CALIN elements in genomes encoding Intl. The CALIN elements might also result from integration of cassettes at secondary sites in the chromosome (9,10,57). We found some cases where IntI was actually encoded in another replicon (3.5% of CALIN). Overall, half of the CALIN were found in genomes encoding IntI and half in genomes lacking this gene.

Insertion sequences (IS) may create CALIN by promoting chromosomal rearrangements in a previously complete integron. The frequency of these events depends on the frequency of IS inside integrons. We therefore searched for IS inside or near CALIN, In0 and complete integrons (see Methods). We found that 12% of CALIN and 23% of the complete integrons encoded at least one IS within their cassettes. Upon IS-mediated rearrangements, the CALIN should be close to an IS. Indeed, 38% of the CALIN had a neighboring IS. Such co-localization was more frequent for CALIN in genomes encoding IntI than in the others (P < 0.001, χ^2^ contingency table). These results are consistent with the hypothesis that IS contribute to disrupt integrons and create CALIN. They may explain the origin of many CALIN elements, especially in the genomes encoding IntI in other locations.

### Divergence of *attC* sites

Since *attC* sites were too poorly conserved in sequence to align using standard sequence alignment methods, we aligned them using the covariance model. We used these alignments to assess the sequence similarity between the R-UCS-L box and the difference in length of VTS sequences between *attC* sites. As expected, both measures showed that *attC* sites were more similar within than between integrons (Figure S9). We then quantified the relationship between the number of *attC* sites in an integron and the average within-integron sequence dissimilarity in *attC* sites. The sequence similarity increased with the number of *attC* sites (Figure 7), *i.e.*, the integrons carrying the longest arrays of cassettes had more homogeneous *attC* sites. Conversely, arrays of heterogeneous *attC* sites were almost always small.

**Figure 7.**
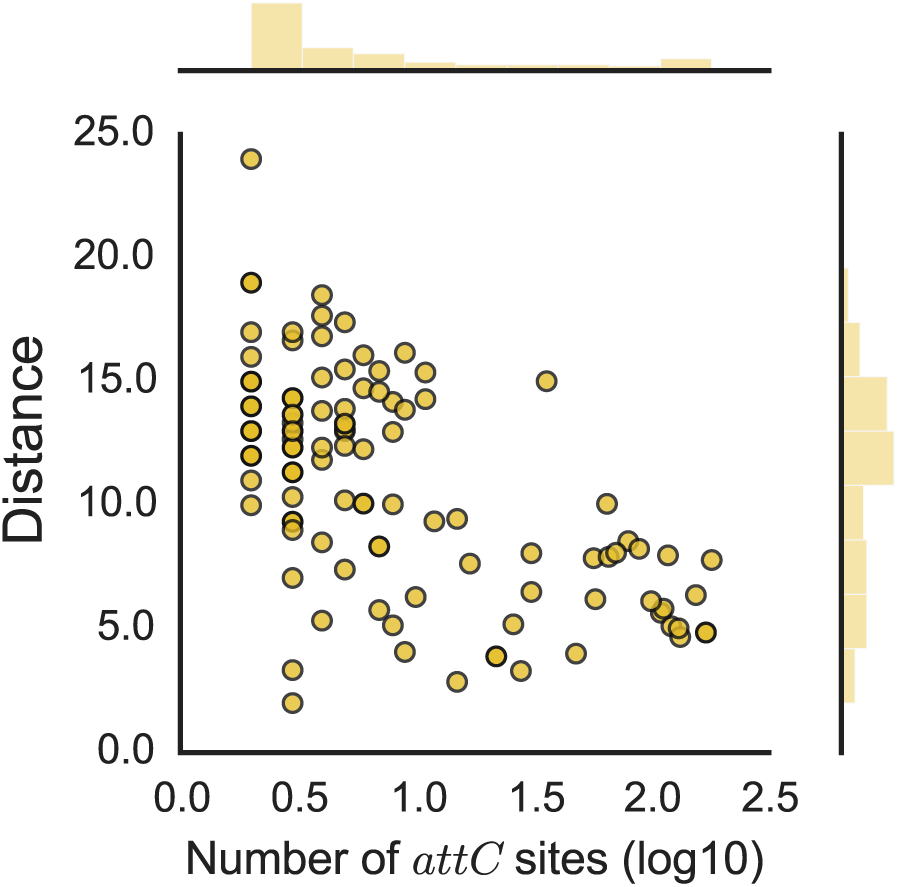
Relationship between the number of *attC* sites in an integron and the mean sequence distance between *attC* sites within an integron. The x–axis is in log10 scale. The association is significant: spearman ρ = −0.53, P < 0.001.

Considering that many previous studies opposed mobile to chromosomal integrons, we tested if our results remained valid when following this dichotomy. We split our dataset into sedentary chromosomal integrons, mobile integrons, and others (unclassified) (see Materials and Methods). Integrons from all three sets were found in the major clades of the IntI phylogeny (Figure S2). Around 67% of the integrons encoded in chromosome were classed as mobile in the species with computed pangenomes (see Materials and Methods), showing that the separation between chromosomal and mobile integrons may be misleading. Expectedly, given their longer arrays of cassettes (Figure 7), the sedentary chromosomal integrons showed more similar *attC* sites than the mobile ones (Figure S9). The similarity of *attC* sites within CALIN elements was between that of sedentary and mobile integron (Figure S9). As proposed before (3), our results suggest that the dichotomy between sedentary chromosomal and mobile integrons may be informative because these two sets are quantitatively different, but may not reflect qualitative biological differences because there seems to be a continuum between large and small integrons.

## Discussion

### IntegronFinder, limitations and perspectives

IntegronFinder identifies the vast majority of known *attC* sites and *intI* genes and is unaffected by genomic G+C content. The high sensitivity with which it identifies individual *attC* sites leads to a very small probability (0.02%) of missing all elements in a cluster of four *attC* sites. Nevertheless, it may be necessary to interpret with care the results of IntegronFinder in certain circumstances. For example, a genome rearrangement that splits an integron in two will result in the identification of a CALIN and an integron (eventually an In0 if the rearrangement takes place near the *attI* site). IntegronFinder accurately identifies these two genetic elements, which are independent from the transcriptional point of view since P_C_ cannot promote expression of the CALIN’s cassettes. On the other hand, these elements may remain functionally linked because cassettes from the CALIN may be excised by the integron-integrase and re-inserted in the integron at its *attI* site. It is unclear if the two elements should be regarded as independent, as it is done by default, or as a single integron. One should note that such cases might be difficult to distinguish from alternative evolutionary *scenarii* involving the loss of the integron-integrase in one of multiple integrons of a genome.

IntegronFinder detects few false positives among integrons and CALIN. Yet, we have identified 431 single *attC* sites in bacterial genomes whose relevance is less clear. Some of these sites might be false positives because their frequency in genomes is close to the upper limit of the false positive rates obtained in our validation procedure. Others might result from the genetic degradation of integron cassettes.

Our study was restricted to the analysis of complete bacterial genomes to avoid the complications of dealing with inaccurate genome assemblies. However, IntegronFinder can be used to analyze draft genomes or metagenomes as long as one is aware of the limitations of the procedure in such data. The difficulty in the analysis of draft genomes results from the presence of contig breaks that often coincide with repeated sequences, such as transposable elements. Their high frequency in integrons implicate that these might be scattered in different contigs. Under these circumstances, IntegronFinder will identify several genetic elements (typically an integron and several CALIN) even if the genome actually encodes one single complete integron. Metagenomics data is even more challenging because it includes numerous small contigs where it is difficult to identify complete integrons. Yet, since the models for *attC* sites and *intI* are very accurate they can be used to identify cassettes and integron-integrases in assembled metagenomes. This might dramatically improve the detection of novel gene cassettes in environmental data.

### Determinants of integron distribution

Our analysis highlighted associations between the frequency of integrons and certain genetic traits. The frequency of CALIN, complete integrons, and In0 is often highly correlated in relation to all of these traits, *e.g.*, all three types of elements show roughly similar distributions among bacterial phyla and in terms of genome size. This association between the three types of elements is most likely caused by their common evolutionary history.

Integrons have well-known roles in the spread of antibiotic resistance. Nevertheless, we identified very few known antibiotic resistance genes in complete integrons outside the class 1 to class 5 integrons. Interestingly, we also found few resistance genes in CALIN elements. This supports previous suggestions that integrons carry a much broader set of adaptive traits, than just antibiotic resistance, in natural populations (28).

We found an under-representation of integrons in both small and large bacterial genomes. Since integrons are gene-capturing platforms, one would expect a positive association between the frequency of integrons and that of horizontal transfer. Accordingly, the lack of integrons in bacteria with small genomes might be caused by the sexual isolation endured by these bacteria, which typically also have few or no transposable elements, plasmids, or phages (58–60). The causes for the low frequency of integrons in the largest genomes must be different, since they are thought to engage in very frequent horizontal transfer (61,62). We can only offer a speculation to explain this puzzling result. Horizontal transfer is often brought by mobile genetic elements. These elements can be very large and costly, while encoding few adaptive traits (if any) (63). The cost of these elements should scale with the inverse of genome size, if larger genomes have fewer constraints on the amount of incoming genetic material and if they select for more frequent horizontal transfer. Hence, the distribution of integrons might result from the combined effect of the frequency of transfer (increasing with genome size) and selection for compact transfer (decreasing with genome size). Further work will be necessary to test this hypothesis.

Most integrons with taxonomic identification available in INTEGRALL are from γ-proteobacteria (90%) (36). Our dataset is more diverse; we found many integrons in β-Proteobacteria and in other large phyla (such as Spirochaetes, Chloroflexi, Chlorobi, or Planctomycetes). This shows that our method identifies integrons in clades distant from γ-proteobacteria. Surprisingly, the genomes from α-Proteobacteria had no integrons, even if they encoded many tyrosine recombinases involved in the integration of a variety of mobile genetic elements. The complete absence of integrons, In0, and CALIN in α-Proteobacteria is extremely puzzling. It cannot solely be ascribed to the frequency of small genomes in certain branches of α-Proteobacteria, since our dataset included 99 genomes larger than 4 Mb in the clade. We also did not find complete integrons in Gram-positive bacteria. It is well known that differences in the translation machinery hinder the expression of transferred genetic information from Proteobacteria to Firmicutes (*e.g.*, due specificities of the protein S1 (64)), but these differences cannot explain the lack of integrons in α-Proteobacteria. Transfer of genetic information between clades of Proteobacteria and between Proteobacteria and Gram-positive bacteria is well documented (65,66). Accordingly, integrons have occasionally been identified in Firmicutes and α-Proteobacteria (36,67), and we found CALIN in Firmicutes and In0 in Actinobacteria. Some unknown mechanism probably hinders the establishment of integrons in these lineages after transfer.

### The evolution of integrons

Our study sheds new light on the evolution of integrons. The use of the covariance model confirmed that *attC* sites are more similar within than between integrons. It also uncovered a positive association between the homogeneity of *attC* sites and the number of cassettes in integrons. If homogeneous *attC* sites result from the creation of cassettes by the integron-integrase, then the largest integrons might be those creating more cassettes.

We found many CALIN in genomes. Previous works have identified *intI* pseudo-genes in bacterial genomes (23,27), and showed IntI-mediated creation of CALIN at secondary integration sites (9,10,57). However, the frequency of CALIN, and especially in genomes lacking integron-integrases, is surprising. These elements may have arisen in several ways. 1) By the unknown mechanism creating novel cassettes if this mechanism does not depend on IntI. 2) By integration of cassettes at secondary integration sites by an integron encoded elsewhere in the genome. 3) By loss of *intI*, even if most CALIN lacked recognizable neighboring pseudogenes of *intI*. 4)By genome rearrangements separating a group of cassettes from the neighborhood of *intI* (as observed in (68)). Mechanisms #3 and #4 are consistent with the presence of IS in a fourth of complete integrons. Mechanisms #1, #2, and #4 might explain why half of the CALIN are in replicons encoding IntI.

There are some similarities, but also key differences, between CALIN and mobile integrons. The average number of cassettes is similar in both elements, but the within-element sequence identity of *attC* sites is different and some few CALIN have many cassettes. CALIN have very few cassettes homologous to those of mobile integrons and far fewer antibiotic resistant genes. This shows that most CALIN do not derive from the class 1 to 5 mobile integrons carrying antibiotic resistance genes.

The lack of an integron-integrase in CALIN elements does not imply that these cassettes cannot be mobilized. We found many co-occurrences of complete integrons, In0, and CALIN in genomes. They might facilitate the exchange of cassettes between elements. Integron-integrases have relaxed sequence similarity requirements to mediate recombination between divergent *attC* sites. It is thus tempting to speculate that integrons transferred into a genome encoding a CALIN might be able to integrate CALIN cassettes in their own cluster of cassettes. Alternatively, CALIN might provide cassettes to naturally transformable bacteria, in case their stable circular forms are able to survive in the environment and be taken up by transformation (69). If integrons often capture cassettes from CALIN elements then many genomes currently lacking integrons but carrying CALIN might be important reservoirs of novel cassettes.

It is unknown whether CALIN genes are expressed or have an adaptive value. Their high abundance in genomes suggests that at least some of them might provide advantageous traits for the host bacterium. CALIN with very degenerate *attC* sites might thus represent an intermediate step between the acquisition of a gene by an integron and its definitive stabilization in the genome by loss of the IntI-based cassette mobilizing activity.

## Availability

The program was written in Python 2.7. It is freely available on a Webserver (http://mobyle.pasteur.fr/cgi-bin/portal.py#forms::integron_finder). The standalone program is distributed under an open-source GPLv3 license and can be downloaded from Github (https://github.com/gem-pasteur/Integron_Finder/) to be run using the command line. Supplementary materials include tables containing all integrons found at different level (elements, integrons, genomes, in Tables S6, S7 and S8). It includes the list of the 596 *attC* sites with their annotated position (Table S2a), and the corresponding file with observed position (Table S2b). We provide the intI_Cterm HMM profile (File S2) and the covariance model for the *attC* site (File S3).

## Acknowledgements

JC is a member of the “Ecole Doctorale Frontière du Vivant (FdV) – Programme Bettencourt”. We thank Didier Mazel, Jose Escudero, Céline Loot, Aleksandra Nivina, Philippe Glaser, Claudine Médigue, Alexandra Moura, and Julian E Davies for fruitful discussions and comments on the manuscript.

## Author contributions

Designed the study: JC EPCR. Made the analysis: JC. Wrote the software and webserver: JC and BN. Contributed with data: MT TJ. Drafted the manuscript: JC EPCR. All authors contributed to the final text of the manuscript.

